# An automated data processing and analysis pipeline for transmembrane proteins in detergent solutions

**DOI:** 10.1101/714303

**Authors:** D. Molodenskiy, H. Mertens, D. Svergun

**Affiliations:** European Molecular Biology Laboratory; EMBL Hamburg; EMBL c/o DESY

## Abstract

The application of small angle X-ray scattering (SAXS) to the structural characterization of transmembrane proteins (MPs) in detergent solutions has become a routine procedure at the most synchrotron BioSAXS beamlines around the world. SAXS provides overall parameters and low resolution shapes of solubilized MPs, but is also meaningfully employed in hybrid modeling procedures that combine scattering data with information provided by high-resolution techniques (*eg.* macromolecular crystallography, nuclear magnetic resonance and cryo-electron microscopy). Structural modeling of MPs from SAXS data is non-trivial, and the necessary computational procedures require further formalization and facilitation. We propose an automated pipeline integrated with the laboratory-information management system ISPyB, aimed at preliminary SAXS analysis and the first-step reconstruction of MPs in detergent solutions, in order to streamline high-throughput studies, especially at synchrotron beamlines. The pipeline queries an ISPyB database for available *a priori* information *via* dedicated services, estimates model-free SAXS parameters and generates preliminary models utilizing either *ab initio*, high-resolution-based, or mixed/hybrid methods. The results of the automated analysis can be inspected online using the standard ISPyB interface and the estimated modeling parameters may be utilized for further in-depth modeling beyond the pipeline. Examples of the pipeline results for the modelling of the tetrameric alpha-helical membrane channel Aquaporin0 and mechanosensitive channel T2, solubilized by n-Dodecyl β-D-maltoside are presented. We demonstrate how the increasing amount *a priori* information improves the model resolution and enables deeper insights into the molecular structure of protein-detergent complexes.

**STATEMENT OF SIGNIFICANCE:** Small angle X-ray scattering (SAXS) using synchrotron radiation is a powerful technique for the structural characterization of transmembrane proteins (MPs) in detergent solutions Overall structural characterization and modeling of MPs from SAXS data is non-trivial, and the necessary computational procedures require further formalization and facilitation. We propose an automated pipeline integrated with the laboratory-information management system ISPyB, aimed at preliminary SAXS analysis and modelling of MPs in detergent solutions, in order to streamline high-throughput studies, especially at synchrotron beamlines.

## INTRODUCTION

Membrane proteins (MPs) are crucial for the normal functioning of an organism as they are involved in essential processes including signalling, nutrient and ion transport, maintaining biological membrane structure and integrity (1–4). They are encoded by about 30% of the human genome (5) and are major targets for modern therapeutic drugs. This makes structural studies of MP solutions extremely important both scientifically and medically (6–10).

The defining structural feature of an integral MP is the presence of a large contiguous apolar surface, facilitating the insertion of the protein into the hydrophobic lipid alkyl chain interior of a biological membrane. In order to characterize MPs *in vitro*, one needs to substitute the natural phospholipid bilayer with an artificial amphiphilic environment that supports the functional state.

At present the majority of investigators studying MPs utilize readily available and inexpensive surfactant systems or detergents (small amphiphilic molecules) (6, 8). However, a common problem observed is the destabilization of protein structure following exposure to detergent molecules (11, 12). Fortunately, the development of milder detergents and detergent stabilized bilayer systems (bicelles) has helped to overcome this problem (7, 8, 13, 14). Importantly, the solution behaviour of many commercially available detergents has been well characterised using a range of techniques, including small angle X-ray scattering (SAXS) (15). Micellar solutions of such detergents have been shown to adopt characteristic sizes, which can be reliably separated from detergent-protein complexes of interest using chromatographic techniques, *eg.* high performance liquid chromatography (HPLC).

Recently, MPs have become increasingly popular objects for small angle scattering investigations (16–19) thanks to the availability of improved online size-exclusion chromatography (SEC-SAXS) set-ups at dedicated SAXS synchrotron beamlines. Integrated HPLC facilitates simultaneous sample purification and X-ray data acquisition, separating the scattering signal from protein, detergent and protein-detergent complexes (PDCs) that appear in the elution profile. Direct measurement of the mobile phase (*ie.* buffer) and elution peaks corresponding to the macromolecule(s) of interest enable a rapid and robust sample characterization following a well-defined sequence of data reduction procedures (20). These procedures are amenable to automation and can be further coupled to advanced analysis and modeling approaches for MP systems, such as those detailed below.

Unambiguous reconstruction of protein shape from a detergent solubilised MP is hampered by the strong scattering signal of bound detergent, typically forming a corona around the lipophilic trans-membrane region. The corona consists of hydrophobic and hydrophilic components, with significantly different scattering/electron densities. Thus the results of standard SAXS modelling procedures are, in such cases, non-realistic and ambiguous. Using complementary data from small-angle neutron scattering (SANS) including contrast matching approaches to isolate scattering from each solution component, the ambiguity of models calculated from detergent solubilised MPs can be reduced (21–24). It was recently shown that SANS data recorded at several contrast points together with an *a priori* estimate of detergent aggregation number, allow one to reliably reconstruct the shapes of MPs in detergent solutions (24). In this publication it was suggested that a minimum of two SANS curves, collected under different contrast conditions, are required to obtain a stable solution. This requirement is necessary to isolate the signals from the two principal contributions to the scattering intensity, from the protein itself and from the hydrophillic part of the detergent corona. For this purpose the author developed an *ab initio* algorithm with a four-phase bead model for a protein: (1) protein, (2) detergent heads, (3) detergent tails, and (4) solvent (24). An additional set of geometrical and symmetrical constraints was also required.

In comparison to shape reconstruction from multi-contrast SANS data, reconstruction of MPs from a single SAXS curve is significantly more challenging as the contributions from different phases are difficult to separate. Nevertheless, in view of the relative ease of access to X-ray facilities and lower sample consumption, compared to that of neutron sources, SAXS-based approaches to MP characterization are increasingly popular. A recent work on the tetrameric water-channel protein Aquaporin0 (25) showed that if an atomic structure of the protein is available, it is possible to reconstruct the entire complex by combining online SEC-SAXS with refractometry measurements. In that case, it was essential to use the available *a priori* information about the PDC in order to provide sufficient restraints on the model for a meaningful reconstruction. In the present paper we further formalize this approach and develop an automatic pipeline to build models of PDC utilizing *a priori* information taken from a dedicated database.

Given the rapidly growing interest in the applications of SAXS to study MPs, systematic storage of the relevant information is urgently required. The ISPyB information system (26), a database dedicated to biological SAXS (BioSAXS) and crystallographic experiments on synchrotron beamlines, offers an appropriate framework. ISPyB is already employed at major European synchrotrons (Petra-III, ESRF, SOLEIL, DIAMOND, MAX IV) to facilitate the data flow beginning with the sample tracking and ending with its structural characterization and visualization of the generated models.

In order to be used in this project, the ISPyB database was extended to store information on the available high-resolution structures of MPs (e.g. homologs or crystallographic models), amino-acid sequences in FASTA format and chemical formulae of the given detergent tails and heads. The latter are used for the calculation of electron densities and approximate sizes of the detergent corona in the modelling. The electron density values can also be provided directly by the user for the cases when the automatic calculations are impossible or the values are not considered to be sufficiently accurate. Depending on the amount of *a priori* information, the pipeline decides whether it is possible to automatically reconstruct the model and, if so, builds it utilizing either the multiphase *ab initio* program MONSA (27) or the hybrid modelling program MEMPROT (28). The starting parameters for the relevant fitting procedures are evaluated based on the available data stored in ISPyB, with the hybrid approach (25) executing upon detection of a high resolution protein component. In the absence of a high resolution protein structure the *ab initio* reconstruction is executed utilizing information about the amino-acid sequence of the MP, chemical formula of the detergent and additional data on the other solvent/solute components to assess the shape of the PDC.

The automated membrane protein pipeline (AMPP) described herein, is implemented at the P12 beamline of the EMBL at Petra-III storage ring (synchrotron DESY, Hamburg, Germany) as an extension of a standard SEC-SAXS pipeline SASFLOW. It is integrated into the beamline control software BECQUEREL (29) operating the beamline. If the MP-related fields in the ISPyB database are lacking, BECQUEREL runs a standard pipeline mode for soluble proteins, using the program DAMMIF (30) for a single-phase *ab initio* shape determination. The MP-related extension starts by default if the connection with ISPyB is established by the control software, and the membrane-specific fields in the ISPyB database are filled. The system has been optimized through extensive testing on different MPs in detergent solutions, and several examples of its applications are presented below.

## MATERIALS AND METHODS

### Materials

The published SAXS data of the tetrameric α-helical membrane channel, Aquaporin0 solubilized by n-Dodecyl β-D-Maltoside (25, 28) was used for the pipeline testing. This sample corresponds to a reliably reconstituted PDC with known geometrical parameters of a detergent corona providing a test case with the modeling program MEMPROT (28). The model of the Aquaporin0-detergent complex (24, 25, 28) was used as a reference for the models generated by AMPP. Additional pipeline tests were conducted using the experimental SEC-SAXS data for the mechanosensitive T2 channel solubilized n-Dodecyl β-D-Maltoside (31) downloaded from the SASBDB database (www.sasbdb.org, ID SASDCY6)) (32).

### Computational methods

The AMPP is implemented as an extension of SASFLOW (33), an automated data reduction and analysis pipeline currently running at P12. AMPP utilizes several modules and modeling programs available in the ATSAS package (34) and, in addition, calls the program MEMPROT (28). The primary data analysis is executed directly upon completion of a SEC-SAXS data acquisition and proceeds in the following workflow (Figure.1):

1. Radial integration of 2D SAXS images by RADAVER (33) (typically 1000-3000 frames are collected for a standard SEC-SAXS run set with one second exposure per image/frame).
2. Determination of the buffer and sample ranges from the elution profile using CHROMIXS (35), an important analysis component of the pipeline specifically designed for SEC-SAXS analysis. CHROMIXS automatically identifies both buffer and sample regions in the elution profile, and performs frame averaging and subtraction to produce the SAXS curve of the purified component for subsequent analysis (Figure 2). It is worth noting that CHROMIXS is capable of identifying multiple elution peaks, providing the users with a set of distinct SAXS curves from individual fractions, if present.
3. Calculation of the overall parameters (radius of gyration *R*_*g*_, maximum size *D*_*max*_, excluded volume *V*_*Porod*_, molecular weight *MW*) and the generation of relevant plots (Guinier plot: ln I(*s*) vs *s*^2^; Kratky plot: I(*s*)*s*^2^ vs *s*; and the real-space distance distribution function: P(*r*) vs *r*) for all subtracted curves. An example of these plots for the mechanosensitive T2 channel data is shown in the generated summary table (Figure.3). These parameters are employed for the evaluation of the search volume to conduct the *ab initio* reconstruction using MONSA or for the hybrid MEMPROT fitting procedures.
4. Querying and collecting the available *a priori* information from the ISPyB database via dedicated Simple Object Access Protocol (SOAP) web services. Depending on the results, one of the three modeling paths is executed for the subsequent modeling (Figure.1):

**Figure 1.**
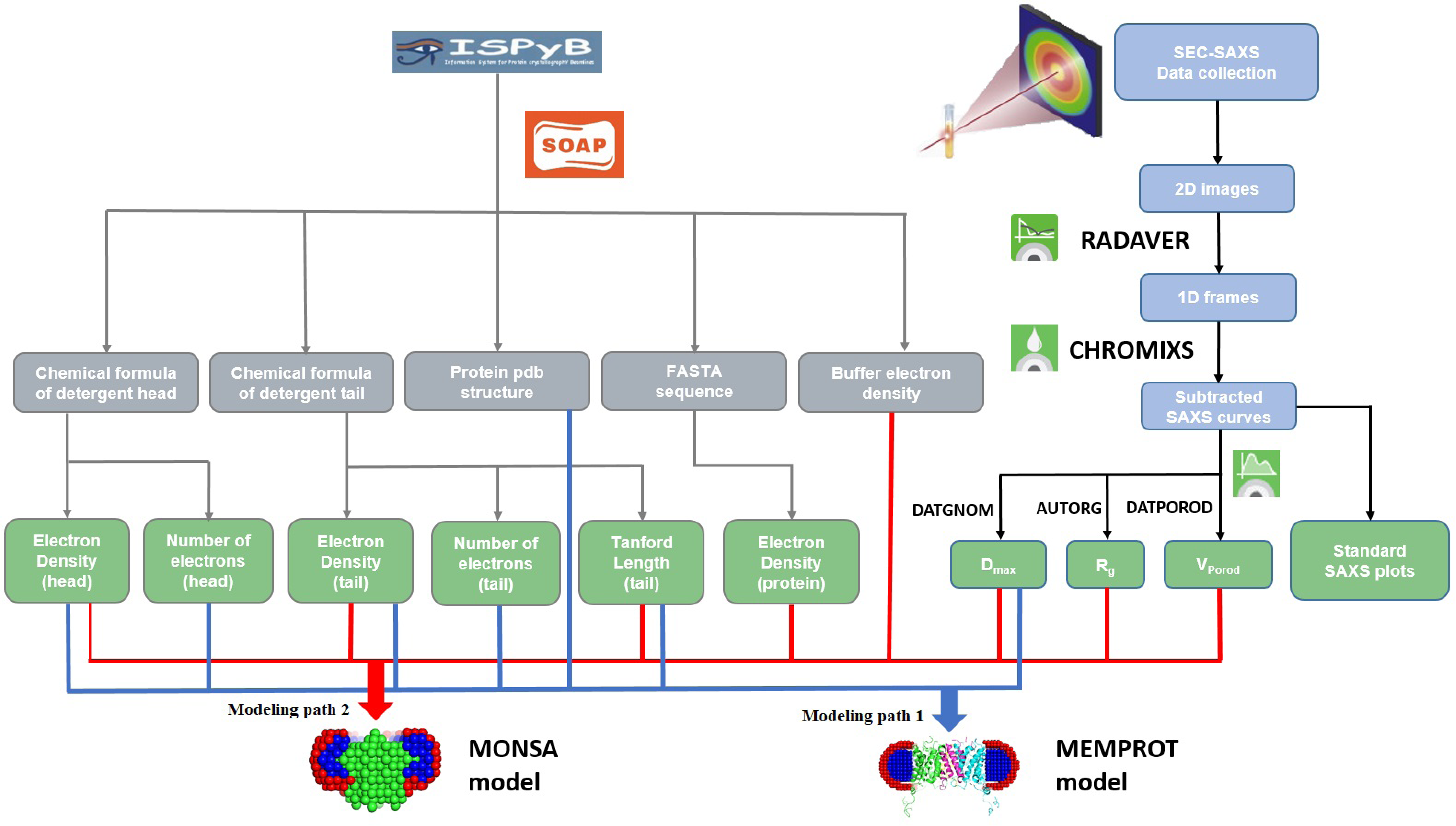
Workflow of the pipeline extension for MPs. The pipeline utilizes *a priori* data from the ISPyB database, calculates overall SAXS parameters and decides, which software to employ for the subsequent analysis.

**Figure 2.**
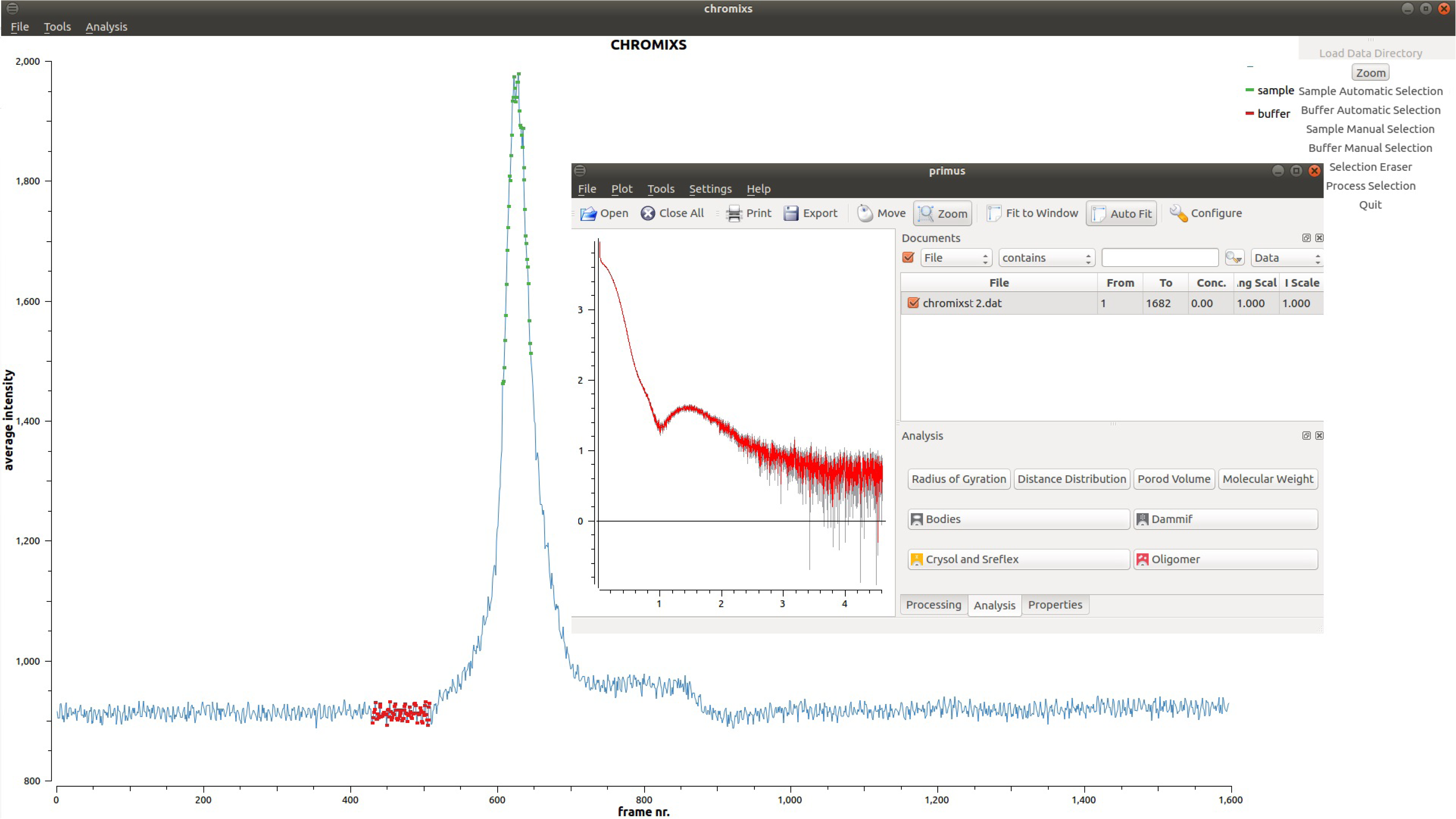
An elution profile of T2 membrane protein, built by CHROMIXS from 1600 consequent SEC-SAXS data sets (average intensity versus frame number). The frames corresponding to the buffer and sample are automatically identified (marked by red and green, respectively), averaged and subtracted one from the other. The resulting curve is displayed in PRIMUS interface (inset).

**Figure 3.**
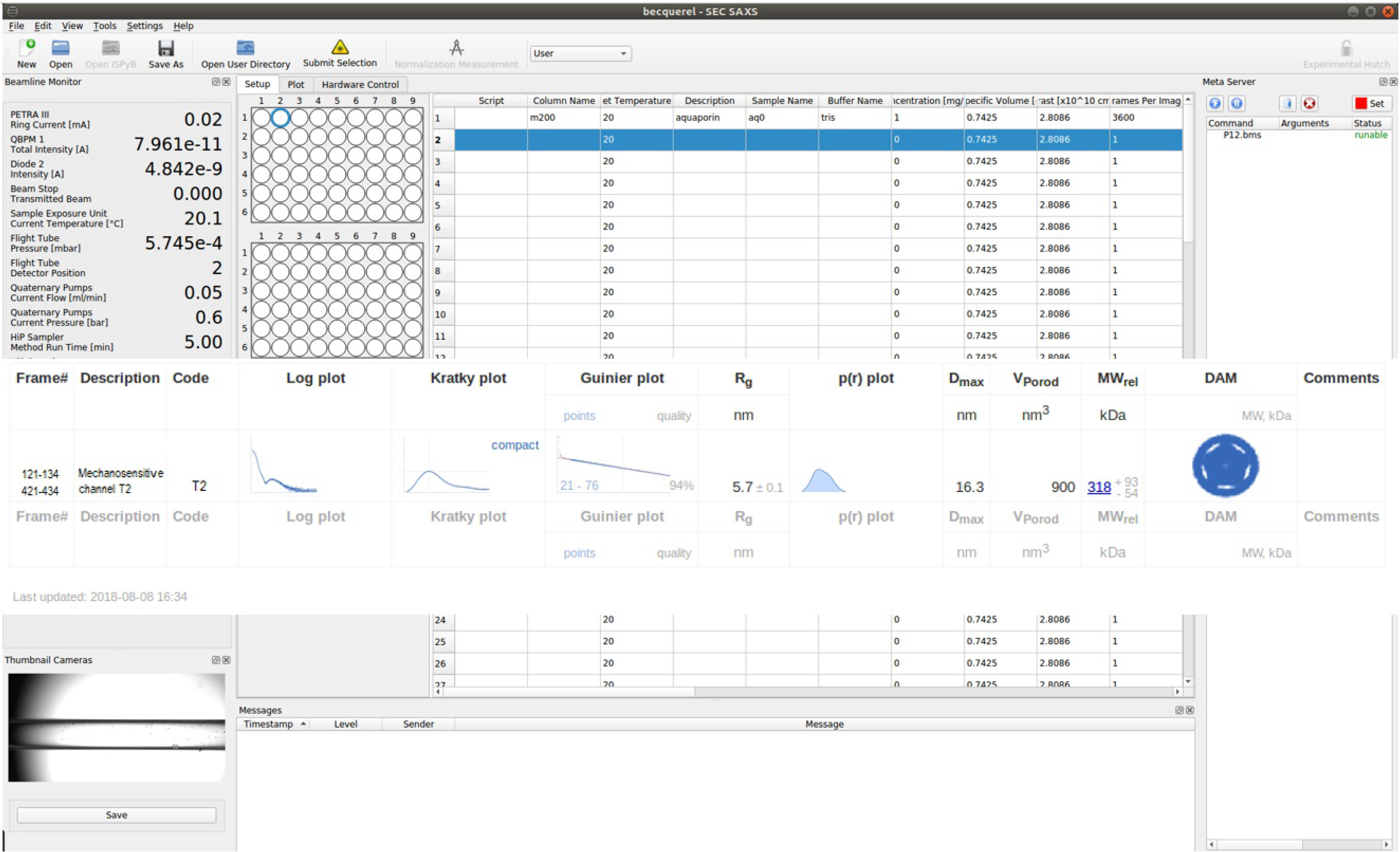
An example of the automatic SAXS analysis of T2 in detergent solution at the P12 beamline (PETRA-III). The sample and buffer ranges of frames on SEC-SAXS chromatogram as determined automatically by CHROMIXS are shown in the **Frame#** field. In this study, a high resolution model was available in the ISPyB database and the modeling was performed by MEMPROT. A thumbnail of the obtained model is displayed in the **DAM** field.

### MODELING PATH 1: High resolution model of the protein and chemical formulae of the detergent heads and tails are available

Initial estimates of the length of the detergent tail and head components are done using the Tanford formula for the detergent tail length: *l*_*c*_ *= 1.5 + 1.265n*_*c*_, where *n*_*c*_ is the total number of carbon atoms. The starting parameters for the evaluation of the corona geometry using MEMPROT (28) are determined as follows:

For a spherical (ellipticity, *e* = 1) detergent torus with the height (*a*) and cross-sectional axes (minor = *b*/*e*, major = *b·e*) equal to the length of a detergent tail (Tanford_tail_), the geometrical parameters are set as: a = b = t = Tanford_tail_ (28) (Figure 4), where *t* is the thickness of the hydrophilic detergent head group. Although the hydrophobic tail of a detergent is typically slightly longer than its hydrophilic head, this assumption appears to be a reasonable first approximation for the further refinement.

**Figure 4.**
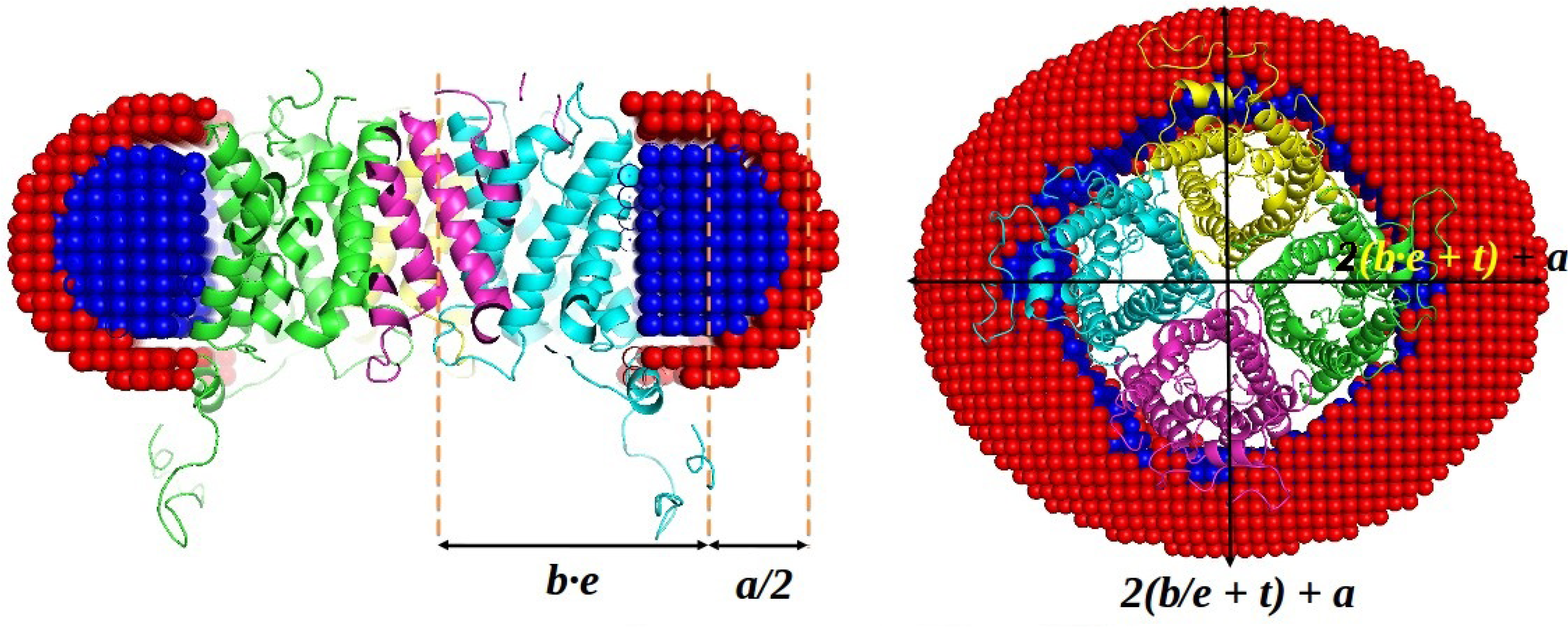
Starting geometrical parameters of a corona for MEMPROT modeling are calculated based on the SAXS curve and *a priori* information from ISPyB. The hydrocarbon tails of the detergent are modelled as an elliptical hollow torus of the height **a** and cross-sectional minor and major axes **b/e** and **be**, where **e** is an ellepticity of the torus.

AMPP employs the MEMPROT minimization procedure, refining the starting parameters within a limited range, estimated empirically based on several tests conducted on different MPs. The a, b and t parameters are searched in the ± 1/3(Tanford_tail_) range, while ellipticity and rotation parameters are kept fixed in order to speed up the computations. The hydrophobic radius of the model is taken as *a + t* = 2·Tanford_tail_, which is in a good agreement with the statistical mean value of transmembrane thickness *D*_*Transmemb*_ ≈ 30 ±3 Å, obtained from the OPM database (36) (https://opm.phar.umich.edu/types/1). The maximum value of *D*_*Transmemb*_ is checked during the modelling such that it does not exceed 45 Å.

### MODELING PATH 2: Electron densities of a protein and chemical formulae of detergent heads/tails are known, but the high resolution model of the protein is not available

The electron densities can be either provided directly *via* the ISPyB interface or calculated from the protein primary sequence (in FASTA format) or from the chemical formula using the built-in look-up tables (37, 38). Knowledge of the detergent chemical formula is mandatory as it provides additional information to constrain the geometry of the MP model. When the high resolution models are not available, *ab initio* low resolution bead modeling is performed by MONSA (27), allowing for three phases with distinct contrasts corresponding to the protein (phase **1**), detergent tails (phase **2**) and detergent heads (phase **3**). For each phase the contrast is calculated and the search space confined to a cylinder and two coaxial spherical hemi-tori (Figure 5A), thus avoiding contacts between the hydrophobic regions and solvent in the final model. A spherical region is fixed for the protein phase in the vicinity of the origin of coordinates, while the rest of the cylindrical region is free to become either protein or solvent. Penalties for the model discontinuity and looseness are applied during the simulated annealing procedure as described in the original paper (27). Three additional boundary regions between the phases (Figure 5A) have a thickness of a few Angstrom allowing MONSA to select the optimum phase assignment during the fitting. The boundary beads (*eg*. between phases **1** and **2)** are allowed to be assigned to either phase **1** or phase **2** (Figure 5A). The model is sought within a spherical search volume of diameter d = *D*_*max*_, where the protein can be surrounded by two-stacked tori (detergent tails and heads) with the lengths evaluated from the Tanford formula (Figure 5B). The transmembrane diameter is calculated as:

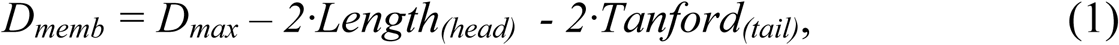

where *Length*_*(head)*_ is considered to be the same as the *Tanford*_*(tail)*_, given that for the most common detergent molecules it is typically smaller or equal to that of the tail. The expected values for toroidal volumes and R_g_ values of detergent phases are calculated on the fly from geometrical considerations (39) and serve as additional constrains while modelling. The total computed volume and *R*_*g*_ of the particle is then compared to the experimental *V*_*Porod*_ and *R*_*g*_, obtained from **step 2**. If available, information from complementary methods (e.g. phase volumes or phase R_g_) can be used to improve the reliability of the final model. Using a spherical 40 beads-per-diameter MONSA grid, together with the above mentioned geometrical restrictions, reliable low resolution models were obtained in a reasonable processing time about 10-15 minutes on a desktop PC (Intel Xeon CPU 3.7 GHz 8 Cores, 12.5 GB RAM). This algorithm is generally applicable to the reconstruction of both transmembrane and membrane-associated proteins. In the case of membrane-associated proteins one can simply shift the fixed “protein” phase to the edge/surface of the detergent disc and extend the protein/solvent phase cylinder outside the entire detergent disc region.

**Figure 5.**
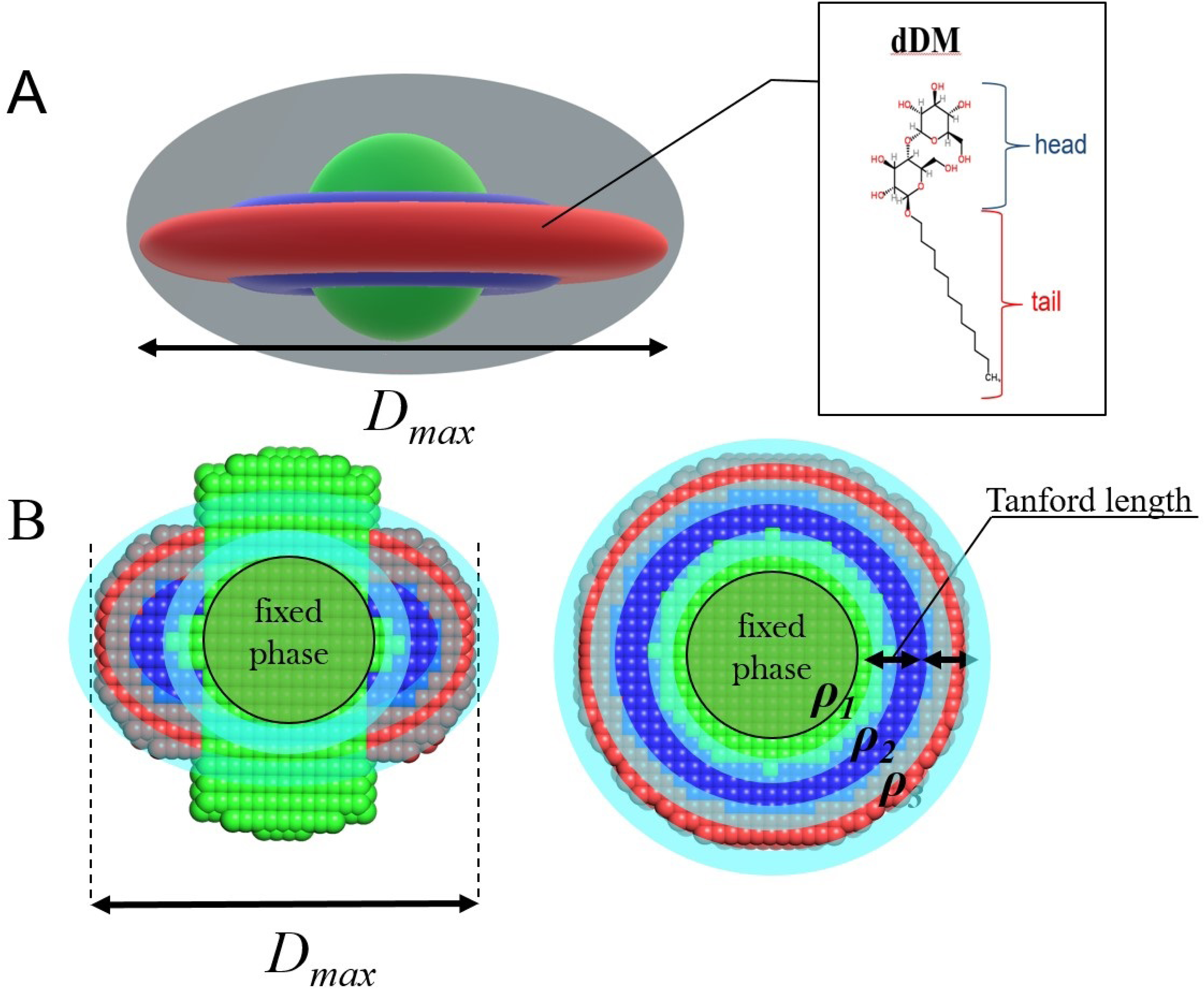
(A) Three-phase grid with boundary regions between phases (marked by cyan) and a model that is sought taking into account phase volumes and radii of gyration (B). D_max_ is considered to be orthogonal to transmembrane region as depicted at the picture. Radius of sphere, exterior and interior radii of tori are calculated from D_max_ and the Tanford length.

### MODELING PATH 3: *A priori* information from ISPyB is not sufficient to generate the required constraints

In this case neither MONSA nor MEMPROT modeling is employed. The pipeline works in the standard SEC-SAXS mode and the low-resolution model is built by DAMMIF (the phase choice only between the PDC and solvent). The pipeline reconstructs low-resolution models from the low-s region (typically *s*_*max*_ < 1.0 nm^-1^) of the experimental SAXS data *ab initio*, minimizing the contribution from the inhomogeneous internal structrure and describing only the overall particle shape. The value of *s*_*max*_ for the initial reconstruction is automatically estimated from the first local minimum of the experimental SAXS curve.

5) As soon as the overall SAXS parameters and the MP model have been computed, the pipeline builds a file in the xml format with the final summary table, gathering together all the results in a compact and human/machine-readable form (Figure 3).

Upon completion the pipeline compresses the calculated 1D SEC-SAXS profiles into a single HDF5 file and reports them back together with the final model into the ISPyB database. The data from the buffer and sample regions, subtracted curves and overall SAXS parameters for each detected peak in the elution profile are displayed in a convenient tabular form (Figure 3). It is worth noting that the summary table generated is applicable to all SEC-SAXS runs conducted at the P12 beamline and is not restricted to the measurement and analysis of MPs. For example, it is often the case that the sample exists in solution as a mixture of different states (*eg.* multiple oligomeric states of a protein or complex), which are separated chromatographically. In this case, AMPP will determine the shapes of individual components in the mixture using **the modeling path 3**. The pipeline is therefore generally applicable.

The results generated can be inspected online using the ISPyB web-server (26). The ISPyB database and web services were extended to accommodate the information on chemical formulae and electron densities of detergents, as well as FASTA amino acid sequences and electron densities of the MPs and buffers. The SEC-SAXS elution profile and the final model can be visualized in the existing ISPyB GUI, as the pipeline output files are compatible with the required format.

## RESULTS AND DISCUSSION

To test the modelling components of the AMPP, we performed an automatic structural characterization of a tetrameric α-helical membrane channel Aquaporin0 in DDM solution. The AMPP was utilized in all three pipeline regimes: i) with the high resolution model available ii) with detergent composition and the protein primary structure (FASTA sequence) known iii) plain *ab initio* modelling (no *a priori* information). The models generated from these three modelling trajectories are displayed in Figure 6 (as the protein was known to be tetrameric, P4 symmetry was utilized in MONSA and DAMMIF).

**Figure 6.**
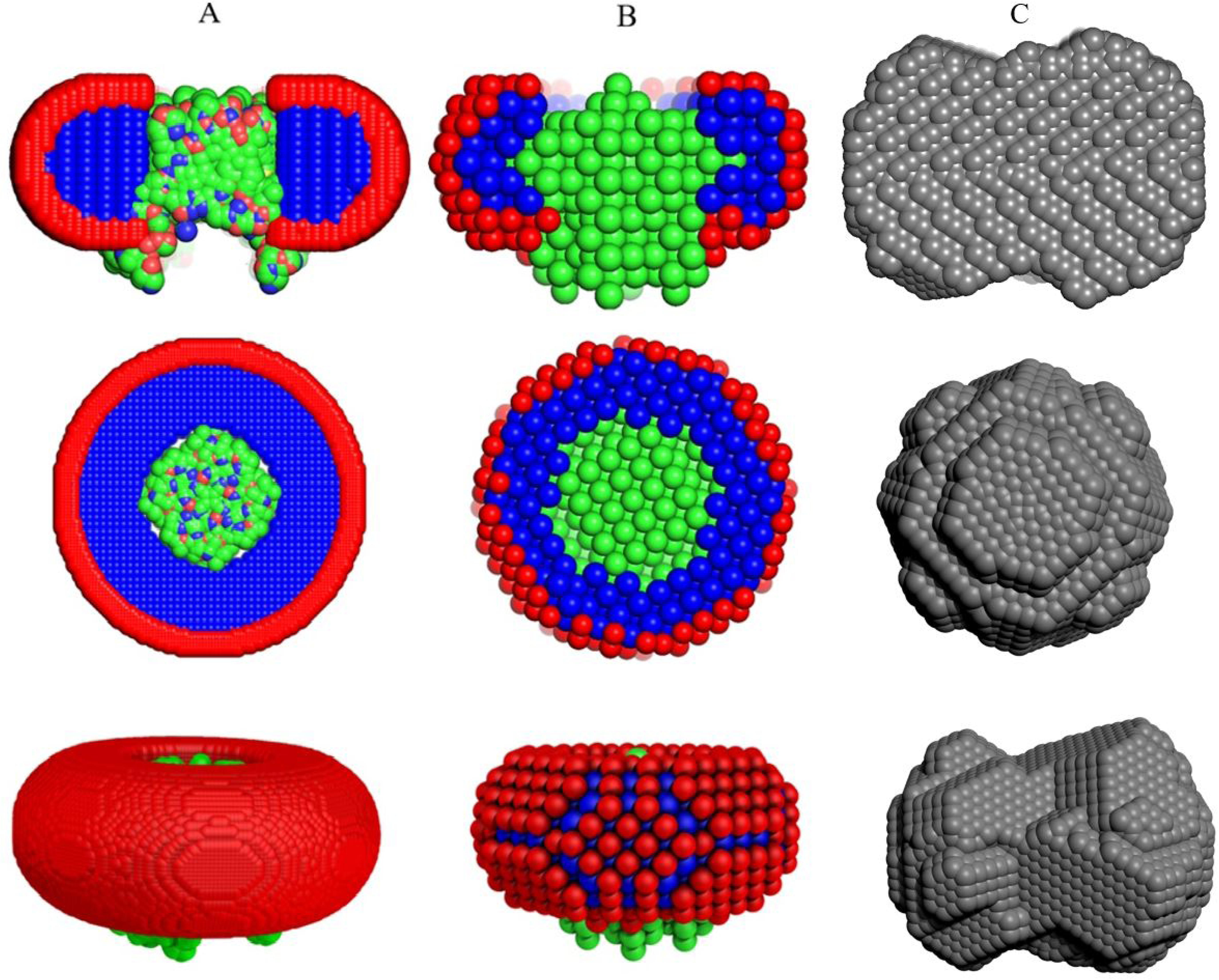
Examples of automatically reconstructed Aquaporin0 PDC models by the AMPP pipeline utilizing MEMPROT (hybrid modeling with known protein model (PDB:2B6P) and unknown detergent belt) (A), MONSA (multi-phase *ab initio*, the model was built from precalculated searching volume; fourfold symmetry around *z*-axis is imposed) (B) and DAMMIF(C) (single phase *ab initio* also with P4 symmetry applied). *The first row* shows the side view cross-section of the PDC components that is represented with a section inside the detergent corona; protein beads are green for *ab initio* models and colored according to atom type for the *hybrid* model, detergent tails and heads are blue and red, respectively. *The second and third rows* represent the top and the side view of the models.

Expectedly, the automatically generated Aquaporin0 models display more adequate structural details with increasing amount of *a priori* information. The single phase based model generated *ab initio* by DAMMIF (Fig. 6C) provides only the overall shape roughly approximating that of the protein-detergent complex. This reflects the limitation of the *ab initio* shape determination procedure, which must assume constant density inside the particle. The more information-rich multi-phase MONSA and MEMPROT models (Fig. 6B, A) display a good level of similarity, with the hybrid model generated using the latter program being more detailed and arguably having a higher resolution. For the multi-phase MONSA modeling path the starting cylindrical search volume used for protein-solvent phases was considerably larger than the final model and contains a spherical core with the fixed protein phase (green area shown in Figure 5B). However, after the minimization procedure the volume occupied by the protein phase becomes only slightly larger than the central fixed spherical volume, and the MONSA models do display protrusions resembling the four α-helical “legs” expected for Aquaporin0. These models are similar to those generated using the hybrid modeling path with MEMPROT, where the protein structure is assumed to be known.

The automatically generated models using the three modeling paths of the pipeline provide good fits to the experimental data at low angles, thus the overall particle shapes coincide and are well described by the data. The deviations observed at higher resolution (Figure 7) reflect contributions from the detergents including the complex contrast situation, which is not accurately captured by the simplified modeling procedures used. This intentional simplification was justified for the sake of speed, such that preliminary models are generated that describe the gross structure of PDCs and do not attempt to describe internal inhomogeneties. One of the simplifications we implemented for the MEMPROT modeling was to assume a horizontal detergent torus with a circular geometry, and a fixed ellipticity *e* = 1. It should be stressed, that further model refinement is recommended in order to obtain more accurate and detailed models, using initial parameters suggested by the pipeline, and as necessary additional penalties as required (such as increased penalty for looseness and/or discontinuity, a higher number of experimental points and dummy beads in the MONSA grid, free ellipticity parameter for MEMPROT, etc).

**Figure 7.**
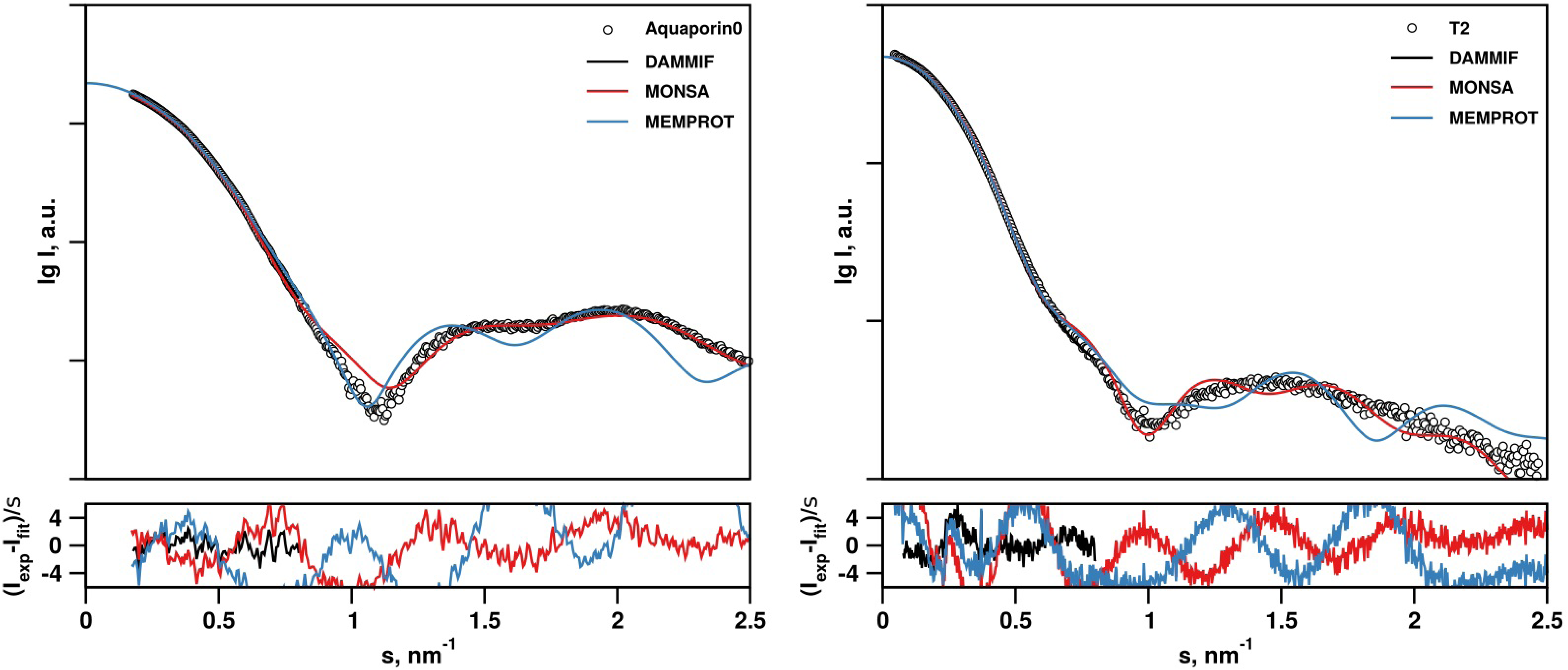
SAXS data from Aquaporin0 (1) and T2 (2) samples and its fit by DAMMIF, MONSA and MEMPROT models. χ^2^ = 1.7 (**1 -** DAMMIF); 12.5 (**1 -** MONSA); 61 (**1 -** MEMPROT); 3.4 (**2 -** DAMMIF); 17.6 (**2 -** MONSA); 20 (**2 -** MEMPROT).

Similar results were obtained from the mechanosensitive channel T2 experimental data obtained at the P12 beamline (Figures 7b-8). There is a high degree of similarity between the three types of models generated, and interestingly, MONSA does reconstruct models with a void in the protein phase (green beads in Figure 8). This void is maintained also when an increased penalty for the bead discontinuity during minimization is introduced. Upon inspection, the high resolution model of a homologous structure, the MscS ion channel of *T. tengcongensis* (PDB: 3T9N) also displays a void in the extracellular domain of the structure. Thus the multiphase *ab initio* reconstruction reliably generates features consistent with those expected for the T2 ion channel.

**Figure 8.**
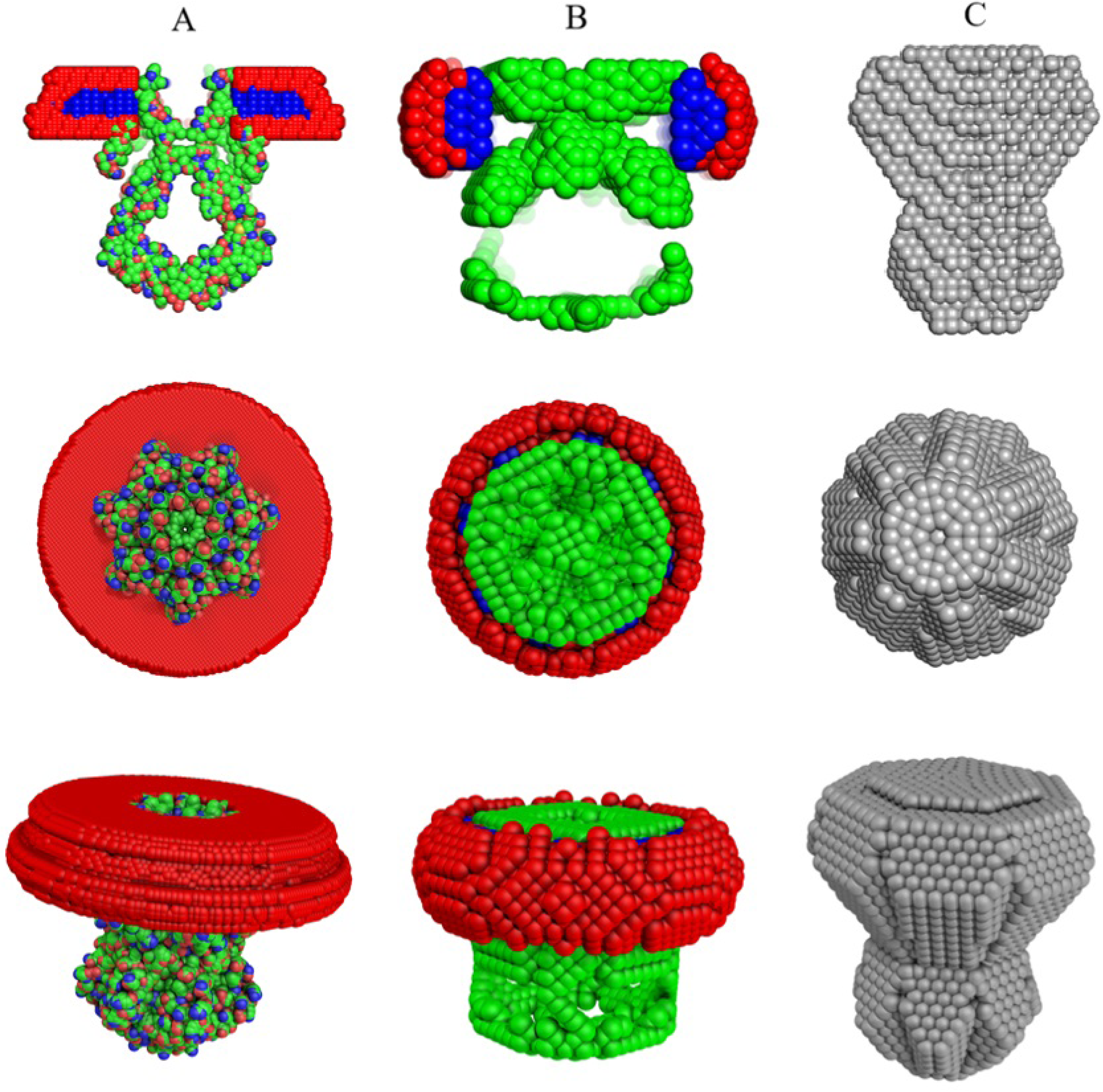
Automatically reconstructed T2 models by the AMPP pipeline utilizing MEMPROT (hybrid modeling: 3T9N pdb model with found shape of detergent corona) (A), MONSA (multi-phase *ab initio* with P7 symmetry) (B) and DAMMIF (single phase *ab initio*, P7 symmetry) (C). *The first row* shows the side view cross-section of the PDC components that is represented with a section inside the detergent corona; protein beads are green for *ab initio* models and colored according to atom type for the *hybrid* model, detergent tails and heads are blue and red, respectively. *The second and third rows* represent the top and the side view of the models.

The reconstructed MONSA model of the T2 channel protein fit the data reasonably well up to s = 0.25 Å^-1^. The on-the-fly generated MEMPROT model fits only up to s = 0.09 Å^-1^, which corresponds to the real-space resolution D = 2π/s ∼ 60 Å.

It is also worth noting that atomistic models are often incomplete, missing *eg.* density for flexible loops which could not be resolved at high resolution. In general, to reliably reconstruct a PDC it is recommended to utilize both methods (full *ab initio* with MONSA, and hybrid with MEMPROT) and compare the results. Thus, the *ab initio* modelling is an important component of the automated MP modelling procedure implemented here.

In summary, an automated SAXS-based data analysis pipeline modified for multi-contrast systems has been developed and implemented at the P12 beamline of the EMBL (Petra-III, Hamburg). The pipeline facilitates the determination of the overall parameters and automated construction of low-resolution structures of solubilized PDCs. Given that the interpretation of the SAXS data from membrane proteins is by far non-trivial task, these results and models are expected to be further refined interactively. However, as demonstrated in the provided examples, AMPP offers at least a good starting point for the subsequent analysis. The relevant data and workflows are integrated into the ISPyB data curation system for SAXS. AMPP is integrated into SASFLOW, an ATSAS pipeline, which is available by request.

## AUTHOR CONTRIBUTIONS

D. Molodenskiy developed and tested the AMPP software. All authors contributed equally to the writing of the manuscript.

## ACKNOWLEDGEMENT

This work was supported by iNEXT, grant number 653706, funded by the Horizon 2020 programme of the European Commission.

